# A low-cost and open-source imaging platform reveals spatiotemporal insight into *Arabidopsis* leaf elongation and movement

**DOI:** 10.1101/2023.08.25.554752

**Authors:** Lisa Oskam, Basten L. Snoek, Chrysoula K. Pantazopoulou, Hans van Veen, Sanne E. A. Matton, Rens Dijkhuizen, Ronald Pierik

**Affiliations:** Plant Environment-Signaling, Dept. Biology, Utrecht University, Padualaan 8, 3584 CH Utrecht, the Netherlands; Theoretical Biology & Bioinformatics, Dept. Biology, Utrecht University, Padualaan 8, 3584 CH Utrecht, The Netherlands; Groningen Institute for Evolutionary Life Sciences, University of Groningen, Nijenborgh 7, 9747 AG Groningen, the Netherlands; Laboratory of Molecular Biology, Wageningen University, 6700 AA Wageningen, the Netherlands

## Abstract

Plant organs move throughout the diurnal cycle, changing leaf and petiole positions to balance light capture, leaf temperature and water loss under dynamic environmental conditions. Upward movement of the petiole, called hyponasty, is one of several traits of the shade avoidance syndrome (SAS). SAS traits are elicited upon perception of vegetation shade signals such as far-red light (FR) and improve light capture in dense vegetation. Monitoring plant movement at a high temporal resolution allows studying functionality, as well as molecular regulation of hyponasty. However, high temporal resolution imaging solutions are often very expensive, making this unavailable to many researchers. Here, we present a modular and low-cost imaging set-up, based on small Raspberry Pi computers, that can track leaf movements and elongation growth with high temporal resolution. We also developed an open-source, semi-automated image analysis pipeline. Using this setup we followed responses to FR enrichment, light intensity and their interactions. Tracking both elongation and angle of petiole, lamina and entire leaf revealed insight into R:FR sensitivities of leaf growth and movement dynamics, and its interactions with background light intensity. We also identified spatial separation in hyponastic response regulation for the petiole and the lamina of the leaf, depending on the light conditions.

## Introduction

Plant life is centred around sunlight, and ensuring access to this vital energy resource is crucial for plants to power photosynthesis. This is particularly challenging in dense vegetation, both in natural and agricultural fields, where plants compete for access to this resource because they mutually shade each other (Casal, 2013; Huber et al., 2021). Upward leaf movement, called hyponasty, elevates the photosynthetically active leaf tips towards higher zones of the vegetation where light availability is improved compared to lower regions. Upward leaf movement is part of the shade avoidance syndrome (SAS) that promotes light capture in dense stands and furthermore includes accelerated elongation growth of stems and petioles, apical dominance and early flowering (Casal, 2013; Pierik & Ballaré, 2021). Interestingly, leaf movements do not just occur in response to competition, but are in fact regulated by a wide variety of cues and environmental conditions. This repositioning can be downwards (epinasty), or upwards (hyponasty). Adaptive relocation of leaves and petioles for example allows plants to align their leaves with the sun to facilitate light capture, or move them away to prevent photodamage (Fu & Ehleringer, 1991; Pérez-Llorca et al., 2019). Leaf angle changes to avert photodamage are further influenced by drought (Pastenes et al., 2005). Upward movement of rosette leaves is also a coping mechanism for high temperature since it facilitates cooling (Koini et al., 2009; Park et al., 2019) and for flooding-escape as it may enable aerial contact (Cox et al., 2003; Lee et al., 2011). Inhibiting leaf movements by physically constraining the leaves was found to negatively affect leaf growth (Cox et al., 2003; Walter et al., 2002). Circadian leaf movements allow diurnal growth and may facilitate overtopping of leaves of neighbouring plants, thus aiding in competition (Woodley of Menie et al., 2019). Overtopping of neighbours is also directly enabled by neighbour-induced hyponasty in response to light cues from neighbours (Bongers et al., 2018; Huber et al., 2021).

Neighbour proximity is perceived by plants through changes in the red (R) to far-red (FR) ratio and blue light levels (Ballaré et al., 1990; de Wit et al., 2016). A low R:FR ratio results from selective FR light reflection by neighbours and can be perceived locally at the leaf tip by phytochrome B (phyB), and result in a hyponastic response of one specific leaf (Pantazopoulou et al., 2017). Neighbour proximity and shade results in upward movement of the petiole and leaf (Michaud et al., 2017; Millenaar et al., 2009; Pantazopoulou et al., 2017; Wit et al., 2012). Upward movement of the petiole is caused by differential elongation between the abaxial and adaxial side (Küpers et al., 2023; Polko et al., 2012). These light-regulated movements are however often studied at relatively low temporal resolution (Pantazopoulou et al., 2017) or very young plants (Michaud et al., 2017), whereas it is known that (subtle) differences in hyponastic response can influence adult plant fitness at the canopy level (Bongers et al., 2018; Pantazopoulou et al., 2021; Woodley of Menie et al., 2019). Altering leaf position can be achieved by adjusting the angle via differential growth rates of the lamina, petiole, or both parts of the leaf simultaneously. The parts of the leaf that can elicit the biggest change in angle are the proximal petiole region (Küpers et al., 2023) and potentially the lamina-petiole junction. Together, this allows for potential independent petiole and lamina angle adaptations for optimal light capture during competition, but such details have not been studied. FR treatments are typically done through discrete FR additions providing strong R:FR ratio reductions, whereas in reality R:FR ratios can fluctuate very broadly. The quantitative relationship between the intensity of the FR signal and the leaf response has not been described. Furthermore, R:FR light variations are often accompanied by fluctuations in light intensity, adding an additional layer of complexity to the regulation of leaf movement that is not well understood. Understanding and further elucidating the mechanisms regulating the hyponastic response requires a high temporal resolution of petiole, lamina and whole leaf movement, as well as a detailed look at the tissues, and their plasticity, that contribute to overall leaf positioning in a canopy.

Measuring petiole and lamina movement and elongation through day and night with a high temporal resolution would aid in further elucidating molecular regulation of leaf movement. Measuring leaf movement was previously achieved with conventional RGB time-lapse imaging (Cox et al., 2003; Millenaar et al., 2005; Rehman et al., 2020). A major disadvantage however is that RGB time-lapse imaging depends on white light, which interferes with photoreceptor activity and night-time imaging in light will likely affect leaf movements. Using infrared lighting and imaging would help solve this potential problem. In some set-ups, leaf lamina and petiole movement are not directly measured, but derived from images from above rather than the side, estimating the leaf angle from variations in projected leaf length (Bours et al., 2012; Rehman et al., 2020). This form of imaging makes it difficult to distinguish whether leaf movement is caused by petiole or lamina movement, and is a derived measure from interpolated growth rates which vary greatly throughout the day. Other high-throughput phenotyping solutions are developed and work well to quantitate leaf movements in great detail. For example, attachable digital inertial measurement unit (IMU) sensors were deployed to measure leaf movement of larger crop species, such as banana, tomato and bell pepper (Geldhof et al., 2021). However, IMU sensors are still too large for use on small leaves such as those of *Arabidopsis*. Alternatively, 3D laser-scanner-based leaf angle imaging methods combined with a robotic stage are successfully used in leaf angle quantifications through time (Dornbusch et al., 2012; Michaud et al., 2017). However, such set-ups are major financial investments and rely on sophisticated downstream data analytics pipelines, making these solutions relatively inaccessible to many research environments around the globe. Another potential draw-back of such set-ups is the relatively modest temporal resolution related to for example potential damage from laser-derived heat dictating that plants cannot be imaged too frequently. Or from measuring multiple individuals using a conveyer belt towards an imaging station, one after the other and thus leaving a relatively long time between repeated measurements on the same individual (Apelt et al., 2015). Additionally, imaging set-ups that depend on a robotic stage, allow for a maximum of one treatment per time, and are restricted by the capacity of the imaging station (Apelt et al., 2015; Dornbusch et al., 2012; Michaud et al., 2017). Thus, taken together, the available methods are often subject to set-up-specific limitations or are too costly to be used by many research groups. There is, therefore, a need for an open-source, low costs solution, such as those enabled by minicomputer based imaging (Tovar et al., 2018).

Here we present a low-cost imaging system with high temporal resolution to accurately follow petiole and lamina movement and growth through time, both during day and night conditions using infrared (IR) LEDs. We also developed an opensource semi-automated image analyses pipeline to analyse petiole, lamina and whole leaf angles and elongation in Arabidopsis. We deployed our set-up to explore how different strengths of R:FR ratio reduction regulate petiole and lamina elongation and hyponasty kinetics. Such FR dose-response data were not previously available and revealed that FR responsiveness scales differently between elongation and hyponastic movement. Additionally, we studied if whole plant FR treatments elicit similar leaf movement and growth kinetics as does localized FR treatment to leaf tips, and how these interact with background light intensity. These data together both show the abilities of the setup and data pipeline, as well as shedding new light on light control of hyponastic leaf movement.

## Results

In response to supplemental FR light that signals neighbour proximity, Arabidopsis plants typically move their leaves upward (Michaud et al., 2017; Pantazopoulou et al., 2017). The response is considered to occur primarily from the petiole base and results from differential elongation between the abaxial versus adaxial side of the proximal petiole region (Küpers et al., 2023). Nevertheless, the angle of the lamina specifically could also change, and the differential elongation for specific lamina movements would then take place in the lamina-petiole junction. To accurately track differential elongation through time in combination with measuring elongation growth, it is essential to monitor the lamina and the petiole separately. Using the images taken in our set-up and the downstream semi-automated image analysis pipeline (Figure 1), petiole, lamina, leaf and junction angles can be measured, together with the lengths of the petiole, lamina and leaf (Figure 2). The leaf tip and lamina-petiole junction are based on user-defined locations in the first image. These are then followed in all subsequent images through Channel and Spatial Reliability Tracking (CRST) based automated tracking of the X-Y coordinates of the moving lamina tip and the lamina-petiole junction using OpenCV (Bradski, 2000; Lukežič et al., 2018; Xu et al., 2019). The information inside a bounding box surrounding the initial user defined locations is combined with information outside the bounding box, using online machine learning to create a discriminative correlation filter (Lukežič et al., 2018; Xu et al., 2019). This approach allows the program to recognise the petiole-lamina junction and the leaf tip in the subsequent images. These changing positions are recorded against the user-defined X-Y coordinates of the petiole origin at the rosette centre (Figure 1C,D) in order to do downstream calculations of lengths and angles. Petiole length and angle are calculated from rosette centre and petiole-lamina junction coordinates, lamina length and angle from petiole-lamina junction and lamina tip coordinates, and leaf angle and length from rosette centre and leaf tip coordinates. From these measurements, several derived components can be quantified, such as the junction angle, the leaf tip elevation and the projected lamina length (figure 2H). The leaf tip elevation and projected lamina length combine the angle with the elongation, showing the total absolute vertical and horizontal displacement of the leaf tip, which helps interpret the plant’s competitiveness.

**Figure 1.**
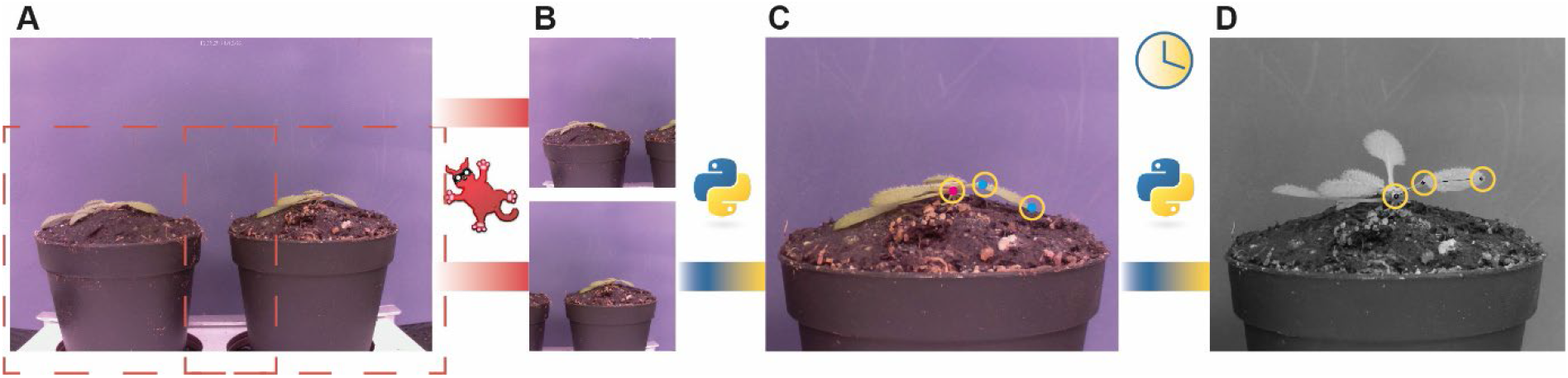
Workflow of the semi-automated imaging pipeline. Example picture taken by Raspberry Pi camera (A), is cropped using Irfan View to create two separate images each containing a single plant (B). With a python script, three locations are user-selected, as indicated by the yellow circles (C). Petiole-lamina junction and leaf tip are selected in the first image (cyan dots). Rosette center is selected on the first and last image (magenta dot). The script will locate these points in all subsequent images (D).

**Figure 2.**
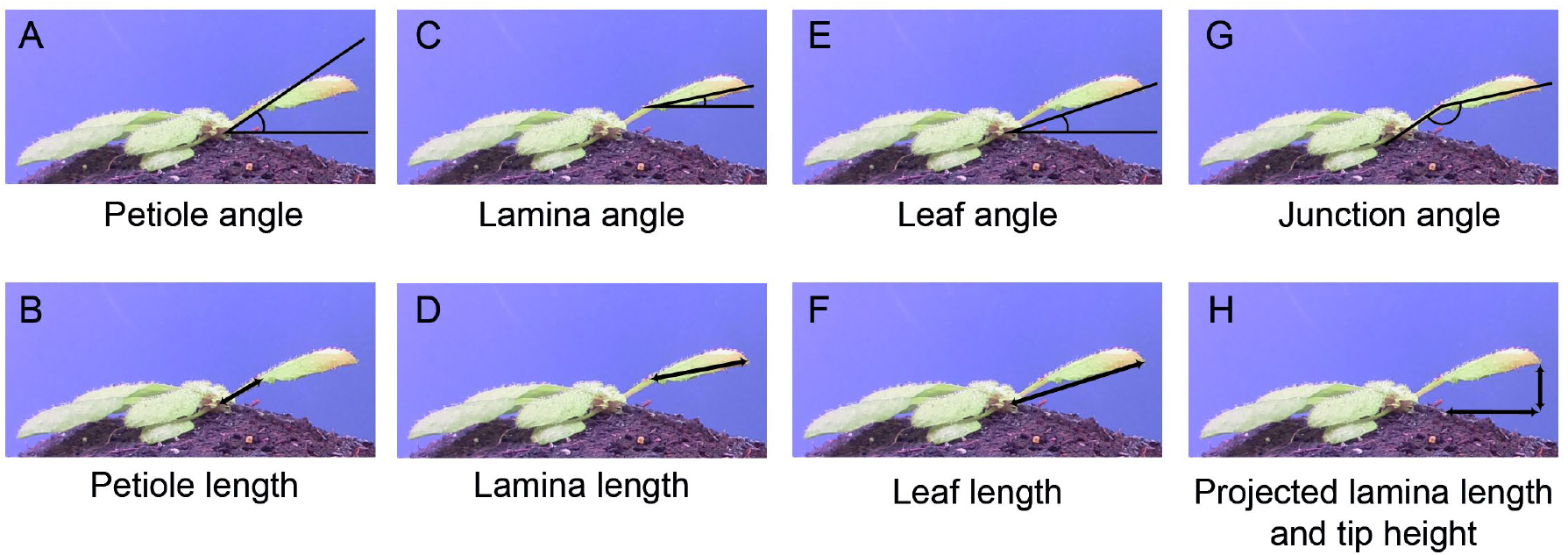
Measurements derived from images taken by the imaging set-up. From rosette heart to petiole-lamina junction: petiole angle (A) and petiole length (E). From petiole-lamina junction to lamina tip: lamina angle (B) and lamina length (F). From rosette heart to lamina tip: leaf angle (C) and leaf length (G). Lamina angle relative to the petiole angle: junction angle (D). Lamina length from petiole-lamina junction to leaf tip based solely on the x-coordinates and lamina tip height based on y-coordinates from the lamina tip and the rosette heart: projected lamina length and tip height (H).

### Subtle differences in timing exist between petiole and lamina movement response to FR light

Using the imaging set-up and analyses pipeline, we explored leaf responses of plants that were placed for 48 hours in either a control white light (WL, R:FR. = 1.5), or WL supplemented with uniform, whole plant FR light (FRw, R:FR = 0.1) (Figure 3). Images were acquired every minute and processed for quantitative data extraction. Control WL plants already move their petioles and laminas in a diurnal pattern (Fig. 3A,B), as previously reported (Michaud et al., 2017; Woodley of Menie et al., 2019). Lamina and petiole patterns are alike in control conditions, but this pattern in movement dynamics between petiole and lamina changes when exposed to a low R:FR ratio. In FRw, the lamina- and leaf angle change peak at the beginning of the night around ZT 12, whereas the petiole angle change reaches the maximum at the end of the night around ZT 22 (Figure 3A, B, C). The junction angle in FRw only differs from WL leading up towards the first night and during the beginning of the first night, starting from ZT 6 until approximately ZT 15 (Supplemental Figure 1A). FRw treated plants display a stronger oscillating pattern than WL indicating rapid angle adjustments and changes in elongation pace (Figure3I and supplemental figure 1B). Oscillation occurs first during the first night, with even more pronounced peaks during the second day between ZT 25 and ZT 28.

**Figure 3.**
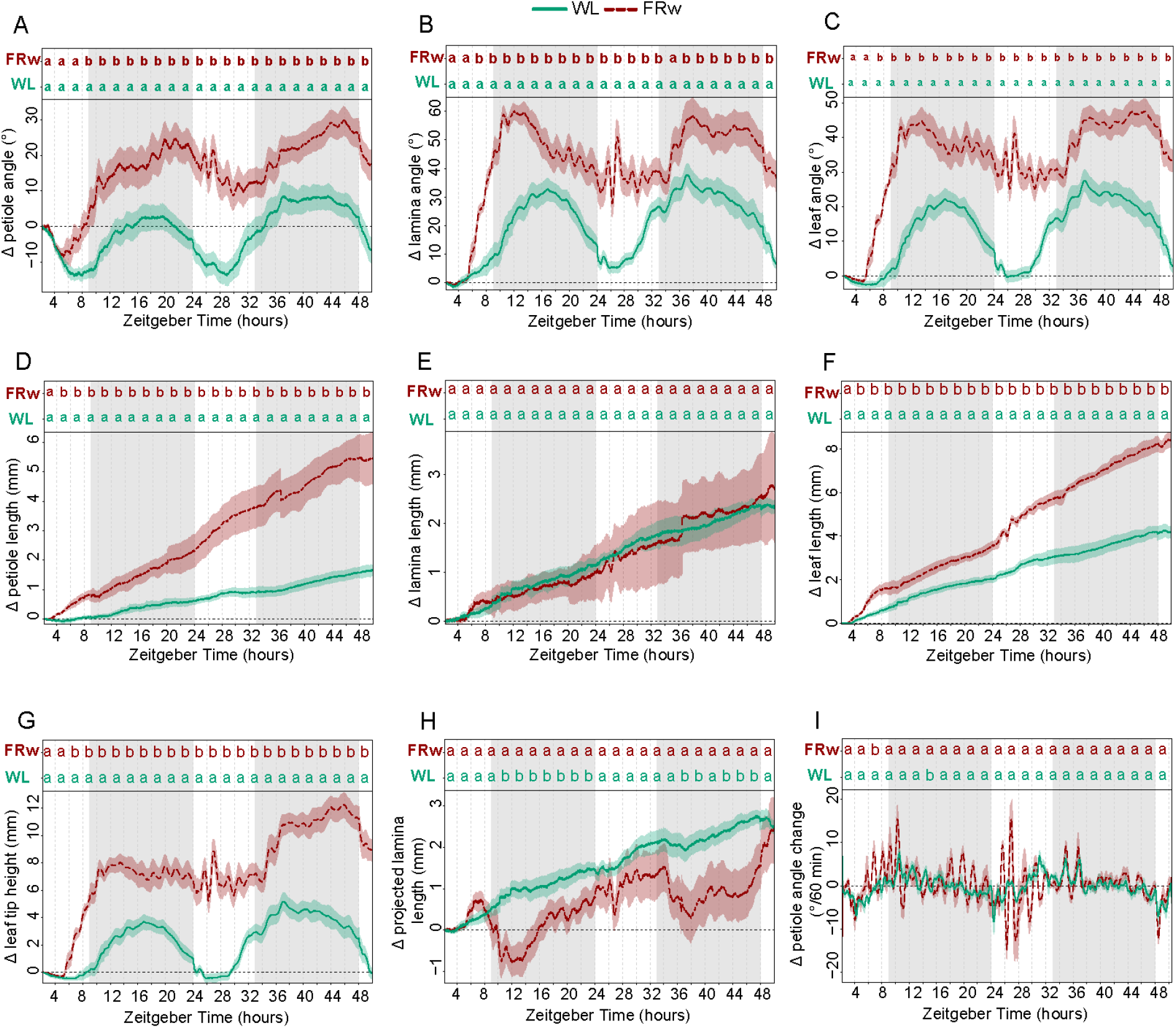
Plant response kinetics to WL or WL with supplemented FR light on the entire plants (FRw). Relative angle change of petiole (A), lamina (B) or total leaf (C). Relative length change of petiole (D), lamina (E) or total leaf (F). Relative leaf tip elevation (G), relative change in projected lamina length (H), speed of petiole angle adjustment over 60 minute periods (I). Grey areas indicate dark period without light or FR exposure. Plants were followed for 48 hours, treated with whole plant supplemental FR (FRw), R:FR = 0.1, or control white light (WL), treatment start time at ZT=2, WL n=7, FRw n= 6. PAR = 140 for both treatments. Letters indicate p < 0.05, calculated per every 2 hours using one-way ANOVA and Tukey post-hoc test.

Petiole and lamina elongation show different FR responses: FRw strongly promotes petiole elongation, whereas lamina length is not affected by FRw (Figure 3D, E, F). Petiole elongation seems to show a difference between FRw and WL slightly earlier than the hyponastic response: around ZT 4 for petiole elongation versus ZT 6 for petiole angle. Leaf tip elevation moves up and lowers again over the course of 24 hours in WL (Figure 33G), but further rises over the second 24 h, driven by both endogenous petiole and lamina movement as well as petiole elongation. These leaf tip elevations are strongly enlarged in the presence of supplemental FR light. Projected lamina elongation is mostly affected during the night for FRw treated plants (Figure 33H), whereas during the day this is not different from WL. Considering the differences in response patterns between the petiole and lamina when exposed to a low R:FR ratio, tracking these organs separately is necessary to accurately monitor plant responses and inform follow-up studies into the molecular regulation of these events that need to be conducted on the appropriate tissues.

### Whole plant FR dosage determines strength of leaf responses

Next, we aimed to investigate the relationship between FR light intensity and leaf responses. We added different fluence rates of FR and maintained a standard background Photosynthetically Active Radiation (PAR: 400 – 700 nm waveband), creating stronger reductions of R:FR ratios with increasing FR intensities. Petiole hyponastic responses are congruent for all four FRw fluence rates during the first hours of the response, but diverge during the night period into two groups (figure 4A) that all have higher petiole angles than the WL control. However, in the second light period, the two higher R:FR ratio’s (i.e. mild FR enrichments) converged with WL 24 hours post treatment start time, whereas R:FR = 0.1 and 0.2 (i.e. strong FR enrichments) maintained their elevated petiole angles. The lamina angle change during the first day and the beginning of the night is increasing proportionally with increasing FR fluence rates, but after 24 hours of treatment, the groups have diverged similarly the petiole hyponastic response (Figure 4B). Junction angle change shows a similar pattern as the lamina angle, except, that the mildest FR treatment (R:FR = 0.6) is barely different from WL control (Figure 4C). Petiole elongation shows an increasing response to increasing FR fluence rates (Figure 4D). Looking more closely at the elongation pattern in the first hours, all four different FRw treatments group together from approximately ZT 4 to ZT 6. Around ZT 8, the petiole elongation of two lowest R:FR ratio groups, FRw 0.1 and 0.2, is significantly higher than that of the two slightly higher R:FR ratio treatments of 0.4 and 0.6. Around ZT 15, the R:FR = 0.4 group show significantly more petiole elongation from the R:FR = 0.6 group, resulting in the gradual increase of petiole length after 24 hours with lowering of R:FR that was previously reported (Bongers et al., 2018).

**Figure 4.**
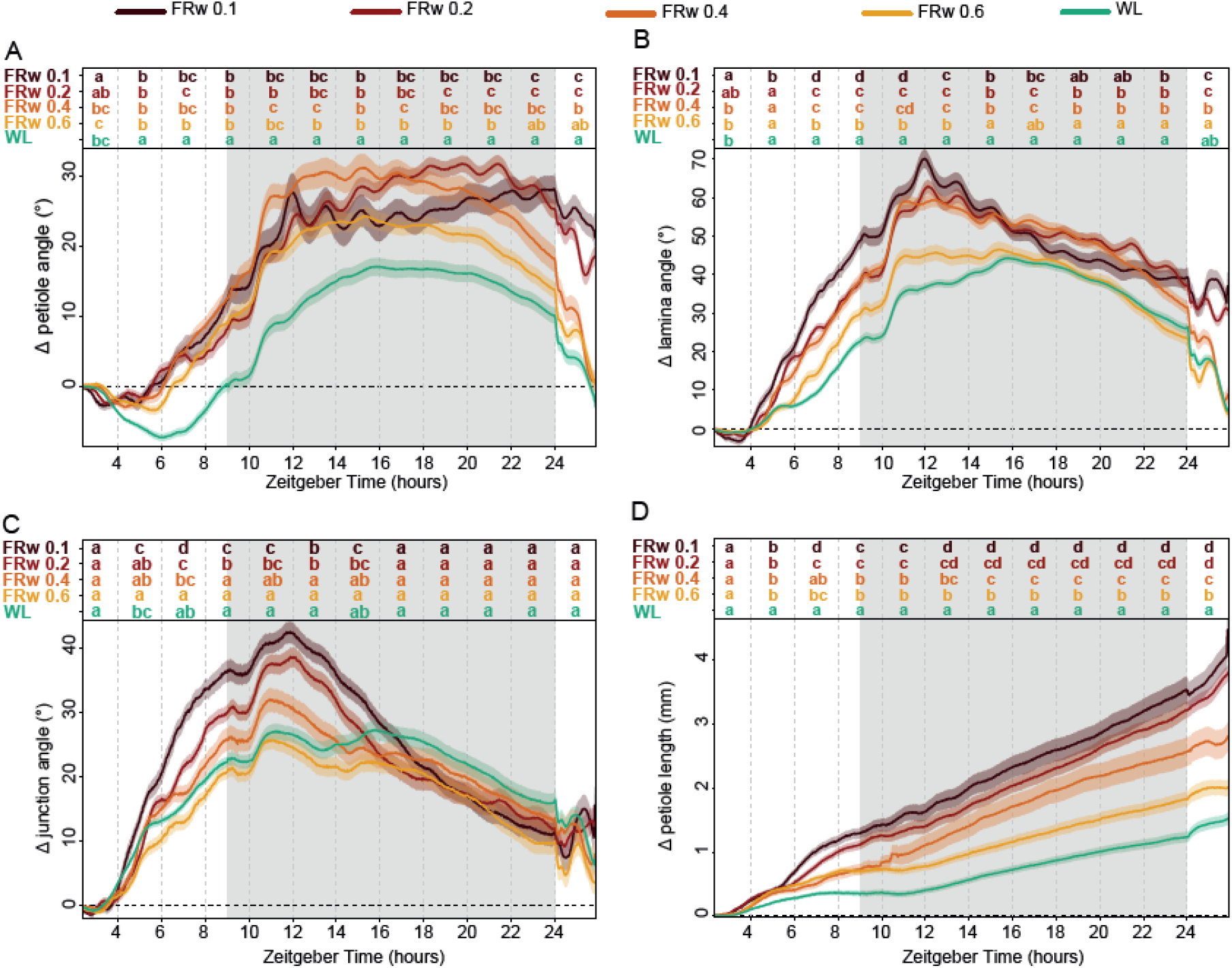
Whole plant supplemental FR intensity influences strength of leaf responses differently between organs. Relative angle change of petiole (A), lamina (B) and petiole-lamina junction (C), and relative length change of petiole (D). Treatments consisted of control (WL) and four FRw doses with R:FR 0.1, 0.2, 0.4 and 0.6 (FRw 0.1, FRw 0.2, FRw 0.4 and FRw 0.6), with 140 PAR for all treatments. Measured for 24 hours with treatment start time at ZT 2, WL n= 69, FRw R:FR 0.1 n = 22, FRw R:FR 0.2 n = 31, FRw R:FR 0.4 n = 28, FRw R:FR 0.6 n = 30. Grey area’s indicate night without light or FR exposure. Letters indicate p < 0.05, calculated per every 2 hours using two-way ANOVA and Tukey post-hoc test.

Increasing supplemental FR fluence rates lead to increasingly strong responses of petiole and lamina growth and movement. However, for the different organs and responses, there are subtle differences: whereas petiole length increase straightforwardly correlates with FR intensity, petiole angle increase is rather similar between all FR fluence rates initially, but the longer-term maintenance of an elevated angle requires a rather high fluence rate of FR light. For the lamina the initial angle increase response is more gradual at the onset compared to the petiole angle, but needs a similarly high fluence rate of FR light to maintain the elevated lamina angle after 24 hours. Together, these results highlight the importance of having both a high temporal resolution and the opportunity to track lamina and petiole separately when assessing the hyponastic response, in order to understand both timing and tissue localisation of leaf positioning.

### Light intensity affects petiole hyponastic response to whole plant FR, but not local FR

Dense canopies not only reflect FR light, but also have reduced light intensities in the lower zones. As PAR and R:FR ratio are not homogenously distributed throughout the canopy, it is important to understand if and how responses to low R:FR are affected by PAR. We therefore moved our adult plants from control conditions of 140 PAR WL to either 40, 80, 140 or 180 PAR. This was combined with supplemental whole plant FR with R:FR = 0.1, where the FR fluence rate was adjusted for every PAR condition such that the FR treatment always reached R:FR = 0.1, irrespective of the PAR combination treatment. We also included a second type of FR treatment: a spotlight FR irradiation to only the tip of the tracked leaf (FRt), leading to a highly localized drop of the R:FR ratio to 0.1 (Figure 5). Local FR treatment on the leaf tip alone is sufficient to elicit a hyponastic response in the petiole of a magnitude similar to, or even higher than, whole plant FR (Supplemental figure 2; Küpers et al., 2023; Michaud et al., 2017; Pantazopoulou et al., 2017). When plants were moved to lower PAR than the control condition, petiole angles increased without additional FR required (Figure 5A, D, G), a well-established response to low light (Millenaar et al., 2009). Surprisingly, at light intensities below 140 PAR, only FR treatment locally on the leaf tip would elicit a hyponastic petiole response additional to the increase seen in the reduced WL fluence rate conditions (Figure 5A, D, G), and whole plant FR had no additional effect. As a separate control, we performed an experiment where we reduced the local FRt fluence rate with the same percentage as used to reduce PAR and found that this lower FRtip fluence rate had the same effect as strong FRt (Supplemental Figure 3). Lamina hyponastic response differed from petiole hyponastic response, since both FRt and FRw induced an additional angle movement in low PAR (Figure 5B, E, H). Higher light intensity (PAR = 180) resulted in a delayed onset of FRw-induced petiole hyponasty compared to PAR = 140, combined with a stronger decline around ZT24 (Figure 5G, J), whereas FRt at PAR = 180 resulted in a similar response as in PAR = 140. Lamina angle changes in PAR = 180 remained similar to control conditions (PAR = 140) for all treatments (Figure 5H, K). Petiole elongation persisted in FRw throughout all PAR conditions, although there is a trend towards milder responses at lower PAR (Figure 5C, F, I, L). These results indicate that there is an interaction between background light fluence rate and the location of low R:FR perception. Furthermore, the petiole and the lamina respond differently to these combinations of background light intensities and low R:FR treatments.

**Figure 5.**
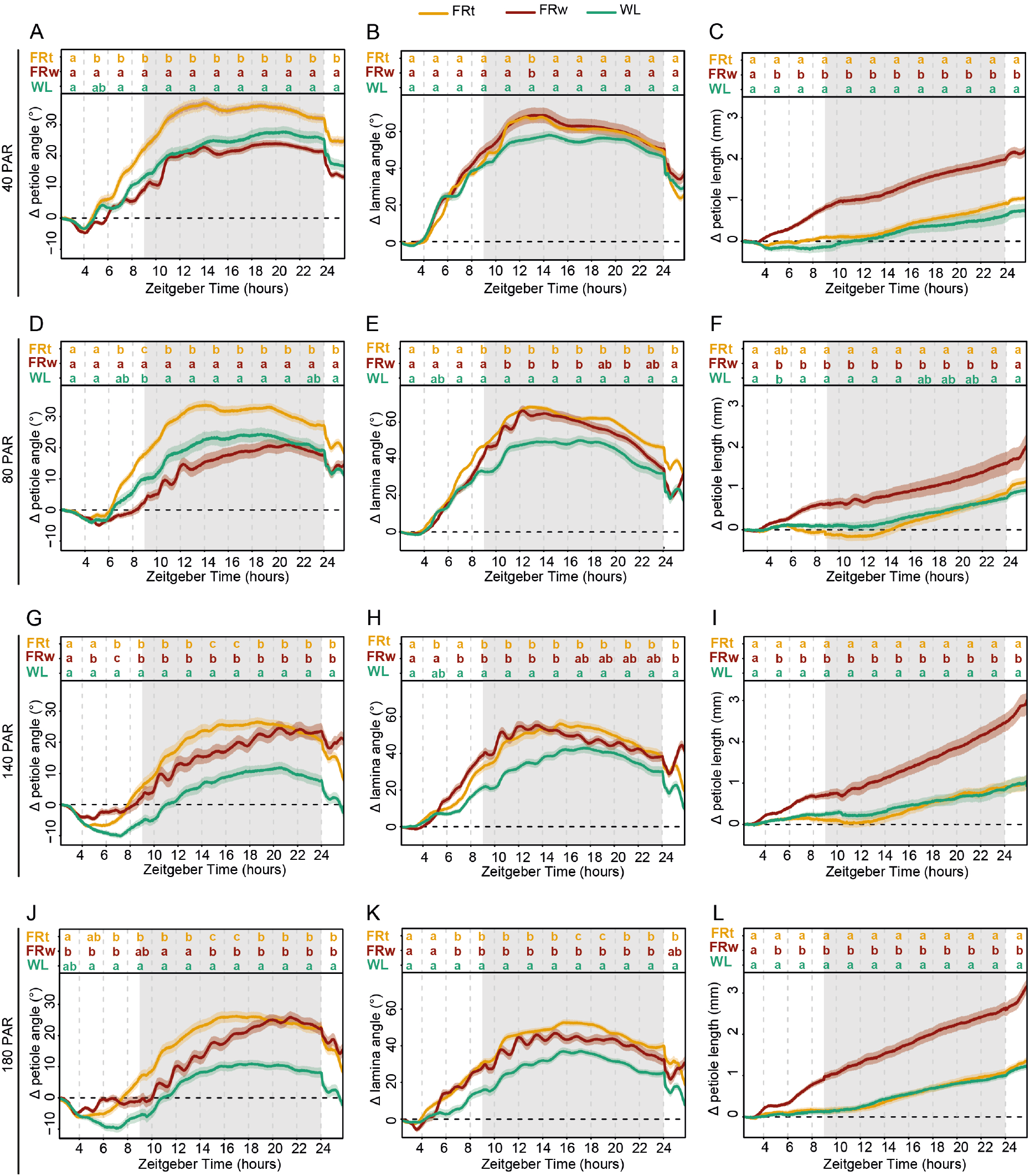
Interaction between background light intensity and local or whole plant supplemental FR treatments. Relative angles of petiole and lamina, and relative petiole elongation with a background light intensty of 40 PAR (A-C), 80 PAR (D-F), 140 PAR (G-I) and 180 PAR (J-L). Treatments consisted of supplemental local FR at the leaf tip (FRt), whole plant supplemental FR (FRw) and white light control (WL). 40 PAR WL n = 12, FRt n = 24, FRw n = 24, 80 PAR WL n = 11, FRt n = 22, FRw n = 27, 140 PAR WL n = 16, FRt n = 19, FRw n = 25, and 180 PAR WL n = 23, FRt n = 22, FRw n = 23. Grey area’s indicate night without light or FR exposure. Measured for 24 hours with treatment start time at ZT 2. Letters indicate p < 0.05, calculated per every 2 hours using two-way ANOVA and Tukey post-hoc test.

## Discussion

We developed a low cost, modular phenotyping set-up to record leaf angle and length dynamics with high temporal precision. Furthermore, we also developed an open-source image analyses pipeline to derive quantitative information from the acquired images. This allowed for refined analyses of leaf movement and elongation for the petiole and lamina separately. Using this new set-up and image analyses pipeline, the petiole and lamina responses to various FR treatments, as a proxy for neighbour signals, were explored. We showed that petiole and lamina angle responses varied not only between different R:FR treatment locations and ratios, but also between the lamina and the petiole itself. These insights highlight both the potential of the set-up and the complexity of plant movement.

### Spatial and temporal resolution from adjustable image acquisition and analysis pipeline

The current imaging set-up is based on raspberry Pi mini computers, however, other companies have produced similar single board computers. Any single computer board that can control a camera and store the images could suffice. Similarly, the camera itself can be replaced by versions with different focal lengths or other characteristics that are tailored to the intended use. This further underscores the adaptability of this system. Furthermore, the set-up is scalable with minimal costs, as the components are relatively inexpensive and widely available.

The subsequent image analyses is relatively straight forward and based on open source software as well . As the points of interest are user-defined by clicking on the image, the script is adaptable as well. The successful imaging under different light treatments during the photoperiod, together with the IR light imaging in the night period, indicate the robustness of the tracking abilities of the script. Some previously established imaging set-ups use images from above (Bours et al., 2012; Rehman et al., 2020), which requires integration of petiole and lamina angle changes with elongation growth into one leaf movement output. Other motion estimation programs are converting the different motions of separate plant organs such as cotyledons and emerging true leaves into one derived relative motion output (Greenham et al., 2015).

In the set-up presented here, a single leaf is imaged from the side, angles and growth of the different leaf components are measured using basic triangulation (Figure 1, Figure 2). There are some drawbacks to the current analysis method. Firstly, petioles and laminas are not always truly a straight line. Any curvature of either the petiole or lamina is currently not taken into account. However, adding more intermediate tracing points to the petiole and lamina would greatly improve the resolution in shape detection and is a relatively straightforward elaboration of the image analysis scripts. Secondly, because the leaf has to be parallel to the camera, only one leaf per plant at a time can be measured accurately. Even though local FR signalling is known not to cause a response in other leaves (Pantazopoulou et al., 2017), whole plant FR certainly does affect several leaves simultaneously, and so do other environmental parameters such as elevated temperature (Koini et al., 2009).

Advantages of the presented set-up on the other hand, are the high spatial and temporal resolution. This allows for the separate analyses of petiole and lamina angles and elongation (Figure 2, Figure 3), and eliminates the need for integration of these separate processes. Secondly, combined and derived measurements of these separate traits provide further insight into the potential benefits under high planting densities. Tip height for instance could be used as an indication for overtopping potential. Projected lamina length combines the angle with elongation to give an indication of the area that is covered by the lamina and could potentially be used as a proxy for its shading surface area potential. Thirdly, with a datapoint every minute, elongation rate and angle change can be measured in great detail (Figure 3I, Supplemental figure 1), which can guide future studies into the molecular mechanisms regulating the different phases of these responses.

### Insights from enhanced temporal resolution

The high resolution of the set-up presented here, provides detailed response kinetics on the elongation and angle changes of both the lamina and the petiole when exposed to a range of different neighbour proximity mimicking signals and (Figure 3, 4,5). Several separate studies have investigated FR light-regulated leaf movement, but typically not with the high spatial and temporal resolution used here (Bongers et al., 2018; Dornbusch et al., 2014; Küpers et al., 2023; Michaud et al., 2017; Pantazopoulou et al., 2017). This comprehensive dataset was therefore not previously available and increases the resolution and depth of existing knowledge. For example, petiole elongation has been previously found to correlate with FR intensity (Bongers et al., 2018). Our results not only corroborate this finding (figure 4D), but also extend it.

We now see that the initial petiole elongation responses for the different whole plant FR intensity treatments are similar to one another, and only diverge after several hours of treatment. This implies that the early initiation of supplemental FR-induced petiole elongation is much more sensitive to modest FR enrichment, than is the longer-term acceleration and consolidation. This might indicate that different molecular regulators drive the early versus later response. Indeed, in very young Arabidopsis seedlings it has been proposed that prolonged FR-induced elongation involves a rewiring of auxin signaling as compared to the early FR-induced auxin response (Pucciariello et al., 2018). Although it is unknown if similar mechanisms regulate petiole elongation responses to FR in adult plants, it does present an example of how different phases of shade avoidance responses involve different molecular mechanisms, and how time-resolved information is key to resolving these mechanisms.

The importance of capturing the kinetics is further highlighted by the lamina and petiole hyponastic responses to the different FR intensities creating the different R:FR ratio’s (Figure 4). If data collection would have occurred only at the start of the experiment (T=0) and after 24 hours (T=24), the conclusions would have been far less subtle. Based on T0 and T24 time points alone, it could be concluded that a dosage with a R:FR = 0.4 and higher is not sufficient to elicit a hyponastic response, only an elongation response (Figure 4A,B,D). However, we showed that all R:FR reductions led to an initial hyponastic response, yet the mild FR treatments converge to white light control conditions 24 hrs after start of treatment. Also here the initial responses could be very sensitive to even mild FR increases, whereas longer-term consolidation and acceleration of the responses involve mechanisms that require stronger FR fluence rates to be activated. Thus, phenotype data interpretation is far more accurate when being able to follow movement through time.

Another interesting observation is that the upward movement of the petiole was preceded by elongation of the petiole (Figure 3I, Supplemental Figure 1B). This difference in timing is consistent with previous observations made on much younger plants under different day/night regimes (Dornbusch et al., 2014), suggesting that, at least in Arabidopsis, FR robustly promotes petiole elongation first, followed by upward movement. However, the opposite order has been described for *Rumex palustris*, a species know to escape flooding using a combination of petiole elongation and hyponasty. Here, petiole elongation will only occur after a petiole angle threshold has been reached via petiole hyponasty (Cox et al., 2003). Future studies could resolve the molecular underpinnings of the differential timing of petiole elongation versus hyponasty in Arabidopsis and other species.

### Insights from spatial resolution

Next to the temporal resolution that permitted close monitoring of the kinetics of plant movement, the resolution in leaf organ tracing enabled us to distinguish nuanced differences between petiole and lamina responses. Under WL conditions, petiole and lamina angle changes and elongation followed a very similar pattern (Figure 3). The different FR treatments, either or not in combination with different light intensities, show that petiole and lamina sometimes respond differently. Combining various background PAR levels and locations of perception of low R:FR treatment further highlighted response differences between lamina and petiole angles (Figure 5). Both local FR and whole plant FR elicited a lamina hyponastic response in the two lower background light treatments (PAR = 40 and PAR = 80). This was different for the petiole hyponastic response at those two lower PARs, where unlike for whole plant FR, local FR on the leaf tip did elicit a response (Figure 5A, D, Supplemental figure 3).

Previous research has shown that an auxin flow from the leaf tip to the petiole base is required and sufficient to cause a hyponastic petiole response (Küpers et al., 2023; Pantazopoulou et al., 2017). It is possible that this leaf tip-induced flow results in a stronger auxin gradient in the petiole compared to whole plant FR, where putative local auxin synthesis on both abaxial and adaxial side of the petiole itself could counterbalance a gradient, yet promote petiole elongation. Previous work by Millenaar et al. (2009) showed the involvement of auxin in low PAR-induced hyponasty. If we assume that low PAR itself also creates an abaxial-adaxial auxin gradient to promote hyponasty (Fig. 5A,D,G), then perhaps the additional effect of FR-tip derived auxin is relatively less, thus creating only mild additional hyponasty.

Additionally, angle responses to whole plant FR exposure revealed a difference in response timing between the lamina and the petiole, with the lamina reaching the highest point at the beginning of the night, versus the end of the night for the petiole (Figure 3A,B). The differences in responses between the petiole and the lamina were further highlighted when exposing the plants to different whole plant FR dosages (Figure 4). Although both petiole and lamina required the same whole plant FR dosage to maintain their elevated angle after 24 hours, the initial lamina angle increase was highest in in the strongest FR treatments, therefore showing a gradual response to the dosage of whole plant FR during the first half of the treatment (Figure 4B). The petiole hyponastic response on the other hand was identical between all four supplemental FR treatments during the first 10 hours of treatment.

These differences in lamina and petiole responses for both elongation and angle could indicate a difference in either molecular regulation, or timing of this regulation, for (differential) elongation between the lamina and the petiole. We found that laminas responded faster to FR treatment. Plus, where the lamina maintained its hyponastic response, regardless of background fluence rates, the petiole exhibited an interaction between background fluence rates and the location of FR perception. The faster response for the lamina could hypothetically be explained by the auxin flow from leaf tip to petiole base that is initiated by FR light (Küpers et al., 2023). This flow will naturally first pass through the lamina, before it reaches the petiole where the differential growth response is driven from its basal-most zone (Küpers et al., 2023). Although the differential elongation for laminas is not described, this would likely happen at the lamina-petiole junction. Further research is needed to establish whether the velocity of the auxin flow or the size of the differential elongation zone is related to the difference in timing between the lamina and the petiole angle responses.

## Conclusion

Taken together, we present a low-cost and modular imaging set-up and analyses pipeline that provides high temporal resolution for both plant movement and growth. Using this setup and including a number of new light treatment comparisons and combinations, we have discussed novel details of tissue-specific responses and the importance of insights into the kinetics to correctly interpret differential leaf elongation responses.

## Materials and Methods

### Plant materials and growth conditions

*Arabidopsis thaliana* Col-0 seeds were stratified at 4°C in the dark for 3 to 6 days on Primasta soil and transplanted to individual 70 mL Primasta-filled pots after approximately 9 days in the light. Plants were kept in WL short day conditions (9 h light, 15 h dark, 70% relative humidity) for 28 days. Light conditions consisted of approximately 140 μmol m^-2^ s^-1^ Photosynthetically Active Radiation (PAR: 400-700 nm), resulting in an estimated phytochrome Photo Stationary State (PSS) value of 0.79 and a R:FR ratio of 1.5 (Supplemental figure 4A) as the default control conditions (Valoya BX-120 modules, NS1 + FR). Plants of 28 days were selected for experiments when the petiole of their 5^th^ youngest leaf was between 4.5 – 5.5 mm long. This leaf was used in all experiments as the focal leaf. Surrounding leaves that blocked view of the camera on the 5^th^ youngest leaf were removed to facilitate down-stream automated image analyses.

### Experimental light and treatment conditions

Plants were grown at 140 μmol m^-2^ s^-1^ PAR and transferred to their respective experimental condition at 28 days. Local FR enrichment was performed using individual L735-06AU LEDs (USHIO) with a peak emission at 735 nm, reducing the R:FR ratio to approximately 0.1 at the sites of application, which was the leaf tip. Whole-plant FR enrichment was achieved with supplemental light from FR-only BX-120 LED bars (Valoya). PAR variations were achieved by adjusting the background light intensity to the desired condition. The treatments consisted of either a range of R:FR ratios, from 0.1 to 0.6, in 140 μmol m^-2^ s^-1^ PAR, or a R:FR ratio of 0.1 under different PAR levels: 40, 80, 140 and 180 μmol m^-2^ s^-1^ PAR conditions respectively (Supplemental figure 4).

### Image acquisition setup

Image acquisition was done using ten Raspberry Pi 4 model B’s (Raspberry Pi Foundation), and ten RPi Cameras model F (Waveshare Electronics), that support infrared (IFR) imaging, shown in Figure 6. We custom-built a Pi mounting plate from and two pots were placed in fixed positions at X distance of the cameras. The setup was fitted with a blue resopal background to facilitate downstream image analyses. Three IFR LEDs TSUS540 (Vishay Semiconductors) were used per plant for imaging during the night with a peak emission wavelength at 950 nm. Images were taken every 60 seconds for 24 hours, and initially stored on a 64GB micro SD card (ScanDisk). Access and control of the Raspberry Pi’s was achieved with a separate computer using a switch (TYPE), a wired private network for the Pi’s via NAT32 IP Router (NAT Software), and VNC Viewer (Real VNC). Data transfer from the Raspberry Pi’s to the separate computer was achieved via open-source software Syncthing (Syncthing Foundation). Time keeping for the Raspberry Pi’s was achieved using Meinberg Network Time Keeping (NTP) package (Meinberg Funkuhren GmbH & Co KG).

**Figure 6.**
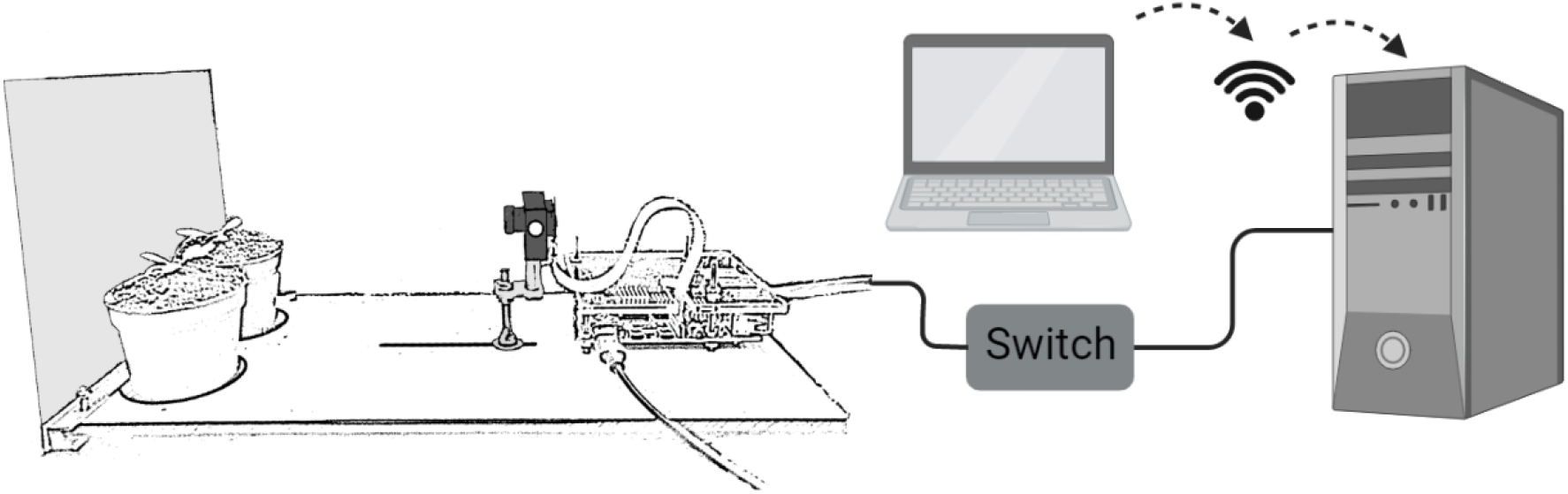
Imaging set-up for leaf angle measurements. Two plants in single pots are placed in front of the Raspberry Pi controlled camera. Ten of these Raspberry Pi and camera combinations are connected to a desktop in a private network via a switch. The desktop can be accessed directly or via a remote desktop connection.

### Image analyses and leaf measurements

Because two plants are imaged per one camera, pictures were cropped using IrfanView 4.58 to result in one plant per image and to reduce computational time in downstream analyses. Cropped images were further analysed in a two-step Python pipeline using two purpose-built scripts, based on OpenCV and ran in Spyder 5.1.5 using Python 3.9.12 The full, open source scripts with descriptions per step are accessible at https://github.com/Pierik-Lab. The first step consists of defining the starting points for the tracking program. Rosette centre, petiole-lamina junction and leaf tip were defined with a locator in the first image, and in the last image (rosette centre). Secondly, a region of known length is selected for downstream conversion from pixels to mm. This information was stored per leaf, and used as basis for the second step of the python pipeline. The second python script performed the actual image analyses and user defined point position tracking. Images were split in R, G and B channels to separate the night time pictures from the light pictures and to enhance the CRST tracking. X and y coordinates of each point of interest were defined per subsequent frame using OpenCV based on information inside and outside the bounding box of 40*60 pixels surrounding the user defined points via an adaptive discriminative correlation filter (Bradski, 2000; Lukežič et al., 2018; Xu et al., 2019). A CSV file per plant was generated containing the x and y coordinates of the points of interest in pixels per frame. Further downstream analyses, angle calculations and graph creation were performed using RStudio 2021.09.1+372 and R 4.1.2 based on the x and y coordinates and is accessible at https://github.com/Pierik-Lab. Petiole, lamina, leaf and junction angles were retrieved via triangulation between meristem, petiole-lamina junction and the leaf tip. The data processing pipeline in R was made such that all data per plant was retained separately, and could be exported grouped in a CSV file. Single plants within an experiment were time synced based on the day-night transition in the pictures. Relative angles and elongation data was obtained by subtraction of the average of the first 5 images from each subsequent image.

### Statistical analyses

Because a datapoint was generated every minute, a grouping of data was done every two hours for statistical testing. One or two-way ANOVAs were performed with R base functions and package emmeans (Lenth, 2022), combined with Tukey tests. Script is available at https://github.com/Pierik-Lab.

## Acknowledgements

The authors would like to thank Otto van de Beek (UU) for technical assistance with the Raspberry Pi hardware set-up and Francis Siefken for help with the IT infrastructure and local network. This work was financially supported by the Netherlands Organisation for Scientific research (NWO) grants Vici 865.17.002 (R.P., L.O.) and ALWOP.509 (C.K.P.).

## Supplemental figures

Supplemental Figure 1. Plant response kinetics to WL or WL with supplemented FR light.

Supplemental Figure 2. Extended timeframe plant response kinetics with WL or WL and supplemental local FR at the leaf tip.

Supplemental Figure 3. Plant response kinetics for reduced supplemental FRt intensity at 80 PAR.

Supplemental Figure 4. Spectral composition of the different light treatments.

