## Supplemental figures for "A low-cost and open-source imaging platform reveals spatiotemporal insight into *Arabidopsis* leaf elongation and movement"

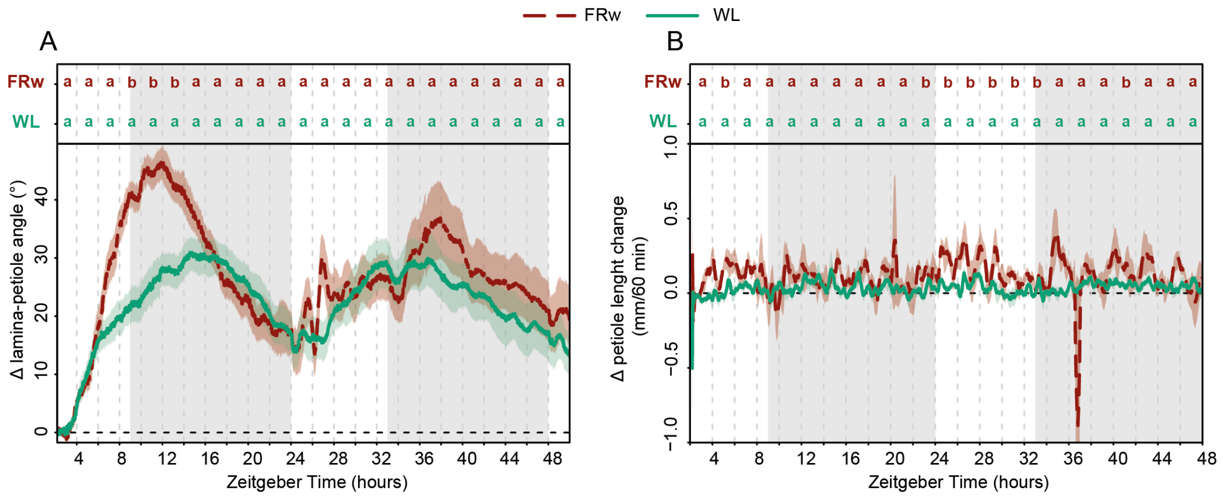


*Supplemental Figure 1. Plant response kinetics to WL or WL with supplemented FR light. Relative angle change for the lamina-petiole junction (A), and speed of petiole elongation over 60 minute periods (B). Grey area's indicate night without light or FR exposure. Plants were followed for 48 hours, treated with whole plant supplemental FR (FRw), R:FR = 0.1, or control white light (WL) with PAR = 140 for both treatments. Treatment start time at ZT=2, WL n=7, FRw n= 6. Letters indicate $p < 0.05$, calculated per every 2 hours using one-way ANOVA and Tukey post-hoc test.*

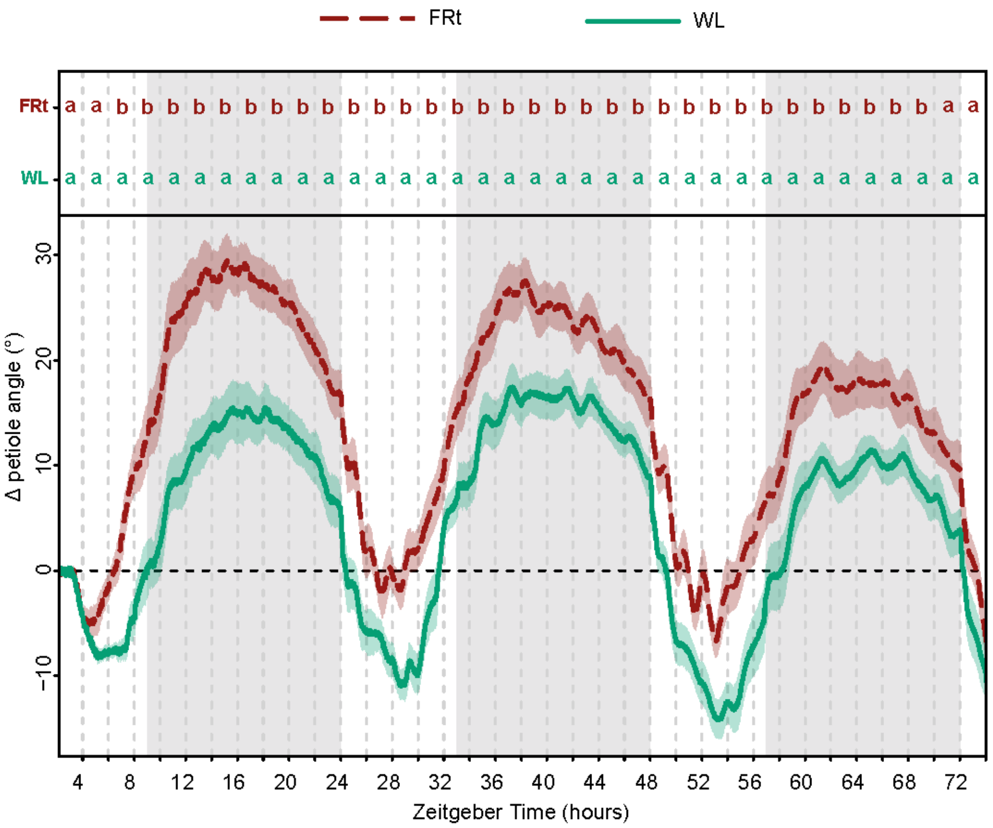


*Supplemental Figure 2. Extended timeframe plant response kinetics with WL or WL and supplemental local FR at the leaf tip. Differential petiole angle over 72 hours is shown for WL and local FR treatment at the leaf tip (FRt) with R:FR = 0.1, and PAR = 140 for both treatments. Grey area's indicate night without light or FR exposure. Treatment start time at ZT=2, WL n=7, FRt n= 8. Letters indicate $p < 0.05$, calculated per every 2 hours using one-way ANOVA and Tukey post-hoc test*

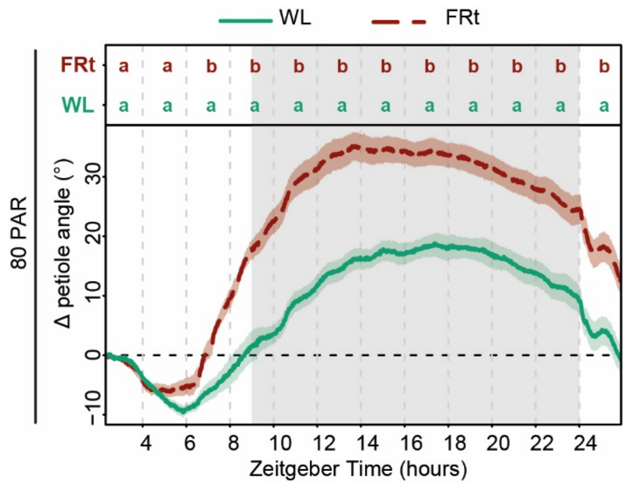


*Supplemental Figure 3. Plant response kinetics for reduced supplemental FRt intensity at 80 PAR. Grey area's indicate night without light or FR exposure. Treatment start time at ZT=2, WL n=8, FRt n= 10. Letters indicate $p < 0.05$, calculated per every 2 hours using one-way ANOVA and Tukey post-hoc test*

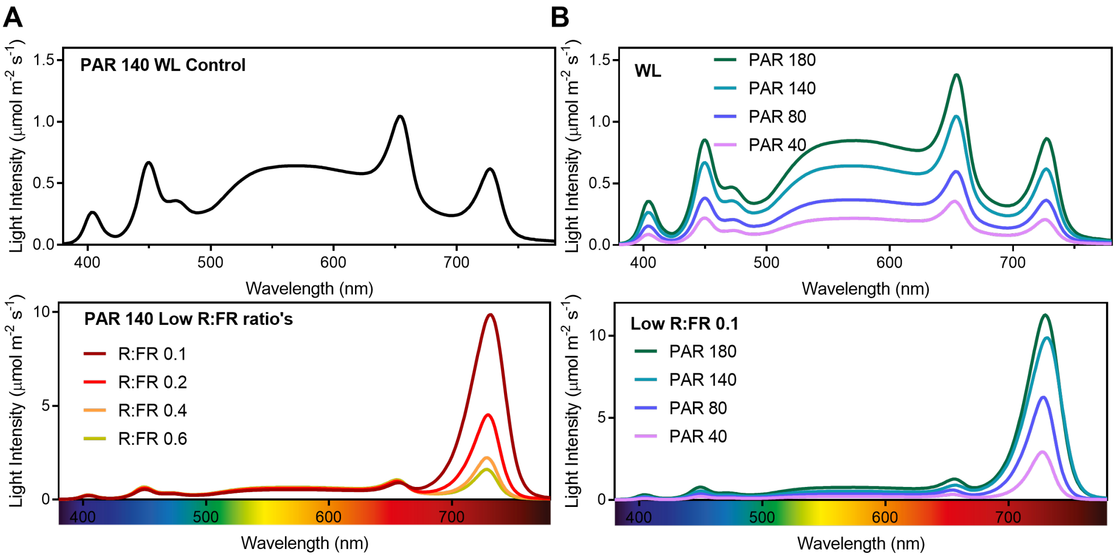


*Supplemental Figure 4. Spectral composition of the different light treatments. Control condition of 140 PAR, with R:FR= 1.5 for the upper panel, and the different FR treatments in the lower panel (A). At the upper panel different WL spectra with R:FR=1.5, together with the corresponding Low R:FR ratio's in the lower panel (B).*
